# Cryptic prophage-encoded small protein DicB protects *Escherichia coli* from phage infection by inhibiting inner membrane receptor proteins

**DOI:** 10.1101/708461

**Authors:** Preethi T. Ragunathan, Carin K. Vanderpool

**Affiliations:** Department of Microbiology, University of Illinois at Urbana-Champaign, Urbana, Illinois, USA

## Abstract

Bacterial genomes harbor cryptic prophages that have lost genes required for induction, excision from host chromosomes, or production of phage progeny. *Escherichia coli* K12 strains contain a cryptic prophage Qin that encodes a small RNA, DicF, and small protein, DicB, that have been implicated in control of bacterial metabolism and cell division. Since DicB and DicF are encoded in the Qin immunity region, we tested whether these gene products could protect the *E. coli* host from bacteriophage infection. Transient expression of the *dicBF* operon yielded cells that were ~100-fold more resistant to infection by λ phage than control cells, and the phenotype was DicB-dependent. DicB specifically inhibited infection by λ and other phages that use ManYZ membrane proteins for cytoplasmic entry of phage DNA. In addition to blocking ManYZ-dependent phage infection, DicB also inhibited the canonical sugar transport activity of ManYZ. Previous studies demonstrated that DicB interacts with MinC, an FtsZ polymerization inhibitor, causing MinC localization to mid-cell and preventing Z ring formation and cell division. In strains producing mutant MinC proteins that do not interact with DicB, both DicB-dependent phenotypes involving ManYZ were lost. These results suggest that DicB is a pleiotropic regulator of bacterial physiology and cell division, and that these effects are mediated by a key molecular interaction with the cell division protein MinC.

**Importance:** Temperate bacteriophages can integrate their genomes into the bacterial host chromosome and exist as prophages whose gene products play key roles in bacterial fitness and interactions with eukaryotic host organisms. Most bacterial chromosomes contain “cryptic” prophages that have lost genes required for production of phage progeny but retain genes of unknown function that may be important for regulating bacterial host physiology. This study provides such an example – where a cryptic prophage-encoded product can perform multiple roles in the bacterial host and influence processes including metabolism, cell division, and susceptibility to phage infection. Further functional characterization of cryptic prophage-encoded functions will shed new light on host-phage interactions and their cellular physiological implications.

## Introduction

Bacteriophages are abundant in the environment with an estimated 10^31^ bacteriophage (phage) particles, and outnumber their bacterial hosts by a factor of 10 to 1 (1, 2). They are found in all ecosystems that harbor bacteria and play a vital role in driving bacterial evolution (3). Based on their lifecycles, phages can be broadly classified as virulent or temperate. Virulent phages use a lytic lifecycle wherein they infect bacterial hosts, use the host cell’s resources to make more phage particles and ultimately lyse the cell to release progeny virions into the environment. Temperate phages can grow using a lytic lifecycle, or alternatively can undergo lysogeny – integrating their genomes at a specific attachment site in the host chromosome and remaining stably associated with the host. The bacterium with an integrated phage genome (prophage) is called a lysogen. Changes in host metabolic conditions or external environmental triggers can induce the prophage, which then excises out of the host chromosome and resumes a lytic lifecycle (4, 5).

Nearly half of all sequenced bacterial genomes have been found to contain at least one prophage, with many genomes containing multiple prophages (6). Lysogeny comes at a cost to the bacterial host due to the extra burden of replication of prophage DNA and the threat of lysogen induction which is lethal to the host cell. On the other hand, there are many well-documented examples of lysogenic conversion, where prophage-encoded products confer new and advantageous characteristics on the host (7, 8). Many prophages carry virulence genes that contribute to the pathogenicity of a bacterial host, *e.g.*, phage-encoded Shiga toxin in *E. coli* O157 strains (9), phage-encoded Diphtheria toxin in *Corynebacterium diphtheriae* (10), and neurotoxin in *Clostridium botulinum* (11). Prophage-encoded toxins, host cell invasion factors and serum resistance proteins promote various aspects of the infection processes carried out by bacterial pathogens (7). Another well-documented benefit of prophages is superinfection immunity. In a mixed population of lysogens and other bacteria, if a prophage becomes induced and lyses a host cell, the active phage particles released will only infect and lyse the non-lysogens, while the lysogens are protected by the prophage-encoded immunity functions (5). Less well-characterized at a mechanistic level are examples of prophage genes that increase the host’s ability to grow under different environmental or stress conditions (12–14).

Growing evidence suggests that in many genomes, most of the resident prophages are cryptic (defective), having suffered mutations that leave them unable to excise from the host chromosome, lyse host cells, or produce infectious phage particles (15–18). A recent study identified and characterized orthologous prophages that were integrated in an ancestral host genome and subsequently passed down vertically with the host chromosome in *E. coli* and *Salmonella* (16). Most of these prophages showed evidence of loss of large portions of the original prophage genome, but the remaining genes were under purifying selection (16). These results suggest that certain prophage genes are selected for during host evolution because they encode products that are advantageous to the host under some condition. The cryptic prophages of *E. coli* K12 have been associated with several host phenotypes, including biofilm formation, stress sensitivity, and antibiotic resistance (19). To understand the molecular basis of cryptic prophage-associated phenotypes, functional characterization of prophage-encoded genes is essential.

In *E. coli* K12, the cryptic prophage Qin carries an operon encoding a small protein, DicB, and small RNA (sRNA), DicF, that both function as cell division inhibitors (20–25). The sRNA DicF represses *ftsZ* translation by directly base pairing with the *ftsZ* mRNA near the Shine Dalgarno sequence (24, 25). DicF also regulates other mRNAs that encode a variety of regulatory and metabolic functions (25). The 62-amino acid protein DicB inhibits cell division by directly interacting with MinC and recruiting it to the septum via interactions with the septal protein ZipA, where MinC stimulates depolymerization of the Z ring, resulting in cell filamentation (23, 26–28). The region immediately upstream of the *dicBF* operon encodes *dicA* and *dicC* and is similar in sequence and structural arrangement to the lambdoid phage immunity locus. DicA is analogous to the P22 phage C2 repressor and DicC to the P22 Cro repressor (29). DicA represses the *dicBF* operon promoter (which is similar to the λ phage P_L_ promoter) and the natural condition(s) leading to induction of the operon are unknown (29). DicB and DicF are conserved in many strains of *E. coli*, and interestingly, many strains of pathogenic *E. coli* possess multiple cryptic prophages encoding *dicBF* operons (25, 30, 31).

In this study, we have identified a role for the *E. coli dicBF* operon in resistance to bacteriophage infection. Short-term expression of the *dicBF* operon promotes *E. coli* resistance to λ phage infection. The resistance phenotype is primarily attributable to DicB. DicB does not affect λ phage adsorption to host cells. Instead, our results suggest that DicB inhibits injection of λ DNA into the cytoplasm through the inner membrane proteins ManYZ, which are components of the mannose phosphotransferase system. Consistent with an effect of DicB on ManYZ activity, we found that growth of *dicB*-expressing cells on minimal media with mannose as the sole carbon source was strongly inhibited. Our results suggest that products encoded by the *dicBF* operon, found in cryptic prophages in many *E. coli* and *Shigella* strains, can impact bacterial physiology, including by altering cells’ susceptibility to bacteriophage infection. We postulate that this may be a common reason why certain cryptic prophage genes are retained in host chromosomes.

## Materials and methods

### Strain construction and media

All strains and phages used in this study are summarized in Table S1 and oligonucleotides (from Integrated DNA Technologies) are listed in Table S2. The strains used in this study are derivatives of *E. coli* K12 strains MG1655 and BW25113. Chromosomal mutations were constructed using λ red recombination method as described in (32–34) or moved into the required strain background using P1 transduction (35).

Construction of strain DB240, which has a P_*lac*_ promoter inserted upstream of *ydfA* replacing the native *dicBF* operon promoter, is described in (25). Oligonucleotides O-PR185 and O-PR186 were used to amplify the kanamycin resistance gene from pKD13 and the PCR product was recombined into the chromosome of DB240 using λ red functions produced by pSIM6 (34). The resulting Δ*dicB*∷kan strain was called PR163. The kanamycin cassette was removed using pCP20 to create Δl*dicB*∷*scar* strain PR165.

A Δ*manXYZ*∷*kan* deletion was moved into DJ624 and DB240 by P1 transduction from YS208 (36) to create PR187 and PR191 respectively. MinC mutants with single amino acid changes E156A and R172A (37) were constructed by first inserting a kan-*araC*-P_BAD_-*ccdB* PCR product in the *minC* gene in strain DJ624. Oligonucleotides O-PR209/O-PR210 (for E156A) and O-PR205/O-PR206 (for R172A) were used to amplify the kan-*ccdB* region of strain YS243 and the PCR product was recombined into DJ624 (pSIM6) to generate strains PR178 and PR179, respectively. Oligonucleotides O-PR211 and O-PR212 (containing the E156A mutation) were used to amplify a segment of DNA from the control strain DJ480 to generate PCR product with the desired mutations for *minC* E156A and recombined into PR178 pSIM6 to generate strain PR180. Oligonucleotides O-PR213 and O-PR214 (containing the R172A mutation) were used to generate a PCR product similarly with mutations for *minC* R172A and recombined into PR179 pSIM6 to create strain PR181. A P_*lac*_ promoter replacing the promoter of the *dicBF* operon, was introduced in PR180 and PR181 to generate strains PR182 and PR183 respectively.

*E. coli* K12 strains were grown in LB medium at 37°C on a rotary shaker. All phage dilutions were made in TM buffer containing 10mM Tris-HCl and 10mM MgSO4, and phage infections were carried out using the same buffer. For phage infections, the top agar was made with equal parts of LB agar and TM buffer, unless specified otherwise (38). Top agar was added to the infection mixture and plated on to LB agar plates.

### Phage propagation

New stocks of each phage were prepared as described in Rotman *et al.* (38). Plating cultures were prepared by growing DJ480 in TB medium with 5 mM MgSO_4_ (and 0.2% maltose exclusively for λ stock preparation) until late log phase, after which an equal amount of TM buffer was added and the mixture was vortexed vigorously. Old phage stocks were titered on the prepared plating culture and plated using top agar, made of equal parts TB agar and TM buffer, onto TB agar plates and incubated overnight at 37°C. The next day, a single individual plaque was punched out and incubated in TM buffer at room temperature for 1-2 hours with occasional vortexing. Between 10 and 30 μl of the single plaque eluate was mixed with 300 μl of DJ480 plating culture and incubated at 37°C for 15 minutes. 3 ml TB/TM top agar was added and plated onto TB plates for incubation at 37°C. After 3 to 7 hours when the lysis was confluent, the plate was overlaid with 5ml TM buffer overnight at room temperature. The TM buffer containing phage was collected in the morning and 4ml fresh TM buffer was added to the plate and kept at room temperature. After 8 hours, the remaining TM containing phage was collected and the combined eluate was centrifuged to pellet the agar and cells down. The supernatant was transferred into a fresh tube, 50ul chloroform was added and the fresh phage lysate was stored at 4°C.

### Efficiency of Plaquing (EOP) assay

The strains used in this experiment were precultured overnight in LB and subcultured in LB medium to ensure all the strains were in the same state of growth when phage infection was carried out. After one hour of sub-culturing (when OD_600_ was ~0.1-0.2), IPTG was added to a final concentration of 0.5 mM to induce the P_*lac*_ promoter. After one hour with IPTG induction, the cells were washed and resuspended in LB medium. Final OD_600_ was measured, and 1 ml of the culture was centrifuged and resuspended in 1 ml TM buffer. 100 μl of phage dilution was added to 100 μl of bacteria from the previous step and incubated for 10 minutes at 37°C. After 10 minutes, prewarmed 3ml LB top agar was added to the mixture and plated onto LB agar plates. The plates were incubated overnight at 37°C and the plaques were counted. EOP was calculated as: (phage titer on test strain in pfu/ml)/(phage titer on control strain in pfu/ml) (39, 40).

### Efficiency of center of infection (ECOI) assay

The strains were prepared for infection as described for EOP assay. The only difference was in the last step of sample preparation, where the cells were resuspended in TM buffer with 0.5 mM IPTG to induce P_*lac*_-*dicBF* during phage infection. The procedure followed for the ECOI assay was based on that described in Moineau, *et al.* (41). λ*vir* lysates were added to 500 μl of prepared strains at a multiplicity of infection (MOI) of 0.1 or less and incubated at 37°C for 10 minutes. The infection mixture was washed with TM buffer containing 0.5 mM IPTG to remove unadsorbed phages and resuspended in 500 μl of fresh buffer. The infected cells were diluted in TM/IPTG buffer and 100 μl of each dilution of strains was added to 100 μl of DJ480 cells in TM buffer, LB/TM top agar was added to this mixture and plated onto LB agar plates. The plates were incubated overnight at 37°C, and the plaques arising from each individual infection were observed and counted. ECOI is calculated as (number centers of infection/ml from test strain) x 100/(number centers of infection/ml from control strain) (41).

### One step growth curve

The samples were prepared for infection as described for ECOI assays. The one-step growth curve experiment was designed based on (41). After resuspending the cells, λ*vir* was added at an MOI of 0.1 or less to 500 μl of cells and incubated for 10 minutes at 37°C. The infection mixture was washed to remove unadsorbed phages and resuspended in 500 μl of TM buffer with 0.5 mM IPTG. The strains were diluted 1:10,000 for DJ480 and 1:1000 for DB240 (P_*lac*_-*dicBF*) to a final volume of 20 ml in LB with 10 mM MgSO_4_ and 0.5 mM IPTG in flasks and incubated in a 37°C water bath. Immediately, 100 μl was withdrawn from the flask and added to 100 μl phage-sensitive DJ480 cells in TM buffer (for lawn formation), added to prewarmed top agar, and plated onto LB agar plates. The first time point was 30 minutes after the start of infection. The same procedure was repeated for each time point. The burst size was calculated as (phage titer at 100 minutes – initial titer at 30 minutes)/ initial titer at 30 minutes. The latent period was calculated as the mid-point of the exponential phase of the growth curve (41).

### Adsorption assay

The procedure described above for ECOI assays was followed and after strains were infected with λ*vir* at an MOI of 0.1 and allowed to adsorb for 10 minutes at 37°C, the strains were centrifuged for 5 minutes at 13,000 rpm to pellet cells and adsorbed phage. 100 μl of the supernatant was removed and dilutions were made in TM buffer. 10 μl of each dilution was added to 100 μl DJ480 (phage-sensitive) cells in TM buffer and incubated for 10 minutes at 37°C. This mixture was plated onto LB plates using top agar and incubated overnight at 37°C. Control titer was calculated using the same procedure as above using the same amount of phage required for MOI of 0.1 added to 500 μl TM buffer (no bacteria). Percentage of adsorption was calculated as (control titer – residual titer)*100/ control titer (39, 42).

### Growth on minimal medium plates with different sugars

For growth assays, M63 minimal medium plates with sugars (glucose, fructose, mannose, N-acetyl glucosamine, and glucosamine) at a final concentration of 0.2% were prepared without or with 0.025 mM IPTG. The strains were streaked on plates and incubated for 44 hours at 37°C. By visual inspection, strains were scored for growth with +++ denoting normal growth, ++ and + denoting decreasing growth and – as no growth.

## Results

### Transient induction of the *dicBF* operon protects against λ phage infection

The region of Qin prophage containing the *dicBF* operon (Fig. 1A) resembles the immunity regions of P22 and other lambdoid phages (29). While functions have not been identified for most of the products of the *dicBF* operon, DicB (a small protein) and DicF (a small RNA) have been shown to inhibit cell division (20, 22–25). We showed previously that DicF post-transcriptionally regulates a variety of genes involved in cell division, growth and metabolism (25). Given their position in the immunity region of the prophage genome, and the fact that the characterized gene products impact cell physiology, we hypothesized that products of the *dicBF* operon could cause changes in the host cell that promote resistance to phage infection. We tested this by comparing phage infections of control and *dicBF*-expressing cells. Since the conditions under which the *dicBF* operon is normally expressed are not fully understood, we used an inducible expression system described previously (25) where we replaced the *dicBF* operon promoter with a P_*lac*_ promoter at the native locus. In addition to the P_*lac*_-*dicBF* strain, we used strains with deletions of different genes in the operon (Fig. 1B).

**Figure 1.**
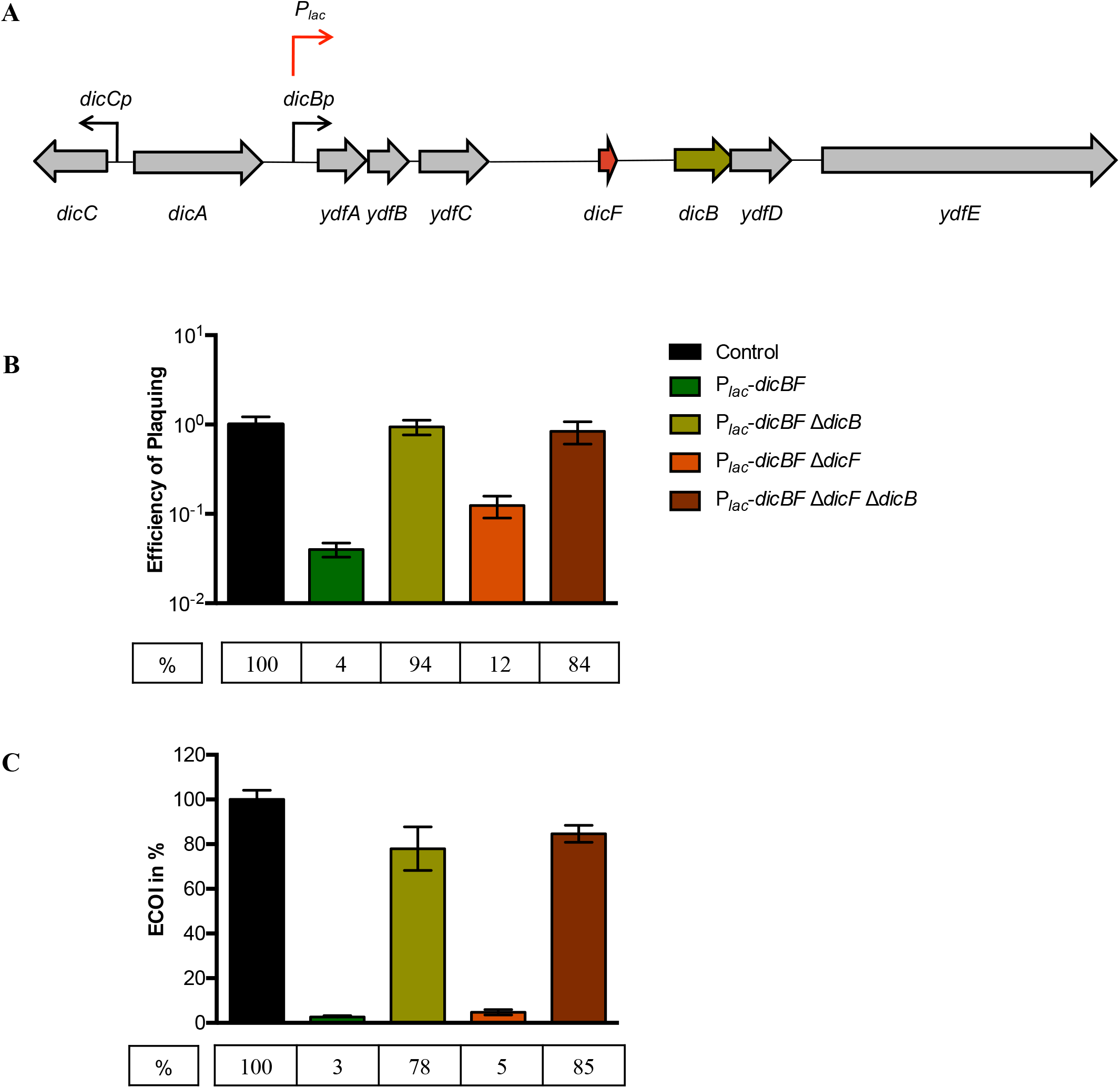
Transient induction of the *dicBF*operon protects against λ*vir* infection. (A) The *dicBF* locus on Qin prophage of *E. coli* K12. The red arrow indicates where P_*lac*_ is inserted on the chromosome, replacing the native *dicBp* promoter. (B) Efficiency of Plaquing (EOP) is defined as (λ*vir* titer on the test strain)/ (λ*vir* titer on the control strain). Strains used in this experiment were: control (DJ480), P*lac*-*dicBF* (DB240), P_*lac*_-*dicBF* Δ*dicB* (PR165), P_*lac*_-*dicBF* Δ*dicF* (DB247) and P_*lac*_-*dicBF* Δ*dicF* Δ*dicB* (DB248). All strains were grown to the same state of growth with the *dicBF* operon induced with 0.5 mM IPTG for 60 minutes. The inducer was washed off, the strains were resuspended in TM buffer, infected with λ*vir* and plated to calculate titer. Error bars were calculated as standard deviation of values from three biological replicates. (C) Efficiency of center of infection (ECOI) is calculated as (number of infectious centers/ml from test strain)* 100/(number of infectious centers/ml from control strain). The strains used in this experiment are control (DJ480), P_*lac*_-*dicBF* (DB240), P_*lac*_-*dicBF* Δ*dicB* (DB243), P_*lac*_-*dicBF* Δ*dicF* (DB247) and P_*lac*_-*dicBF* Δ*dicF* Δ*dicB* (DB248). The cells were grown with induction of the *dicBF* operon with 0.5 mM IPTG and infected with λ*vir* at an MOI of 0.1. The unadsorbed phages were removed, the phage-host complex was added to phage sensitive cells (DJ480) and plated onto LB agar for counting plaques (infectious centers). Error bars were calculated as standard deviation of values from three biological replicates.

We measured phage infection of these strains by Efficiency of Plaquing (EOP) assays, initially using phage λ. In this assay, the phage is titered on all bacterial strains and the titer in plaque forming units (pfu)/ml is calculated for each strain. The EOP is defined as: phage titer on test strain/phage titer on control strain. The control strain lacked the P_*lac*_ promoter. Strains with the P_*lac*_ promoter driving *dicBF* expression were exposed to IPTG for 60 minutes. Then, strains were infected with λ*vir* (a λ mutant that can only complete the lytic cycle during infection of host cells) and plated for titer as described in Materials and Methods. The EOP for the P_*lac*_-*dicBF* strain was 0.04 (Fig. 1B), meaning that rate of infection of the *dicBF*-expressing strain was only 4% relative to the control strain. This result suggested that transient expression of the *dicBF* operon conferred resistance to infection by λ*vir*. To further characterize the basis for this phenotype, we deleted *dicF* and *dicB*, singly and in combination because previous studies identified growth or cell division phenotypes associated with these genes (23, 25). Phenotypes of deletion mutants demonstrated that *dicB* played the most prominent role in the resistance phenotype (Fig. 1B). Deletion of *dicB* alone or *dicB* in combination with *dicF* restored the EOP of λ*vir* to nearly that of control. In contrast, deletions of *dicF* alone had a minimal effect on the resistance phenotype (Fig. 1B). We also carried out infections with wild-type λ phage, and saw similar results for EOP on control, *dicBF*-expressing and deletion mutant strains (Fig. S1).

Because previous studies showed that ectopic expression of the *dicBF* operon impairs growth of the host strain, we reasoned that poor growth of test strains could influence the results of EOP assays. To more accurately assess the outcome of a phage infection on cells expressing the *dicBF* operon, we conducted center of infection (COI) assays. For this assay, strains (Fig. 1B) were induced with 0.5 mM IPTG and λ*vir* infection was carried out at a multiplicity of infection (MOI) of 0.1. After adsorption of phage to test strains, the unadsorbed phages were removed by washing and the infected test cells were diluted and mixed with the phage-sensitive control strain. Productive infections of the test strain are detected as plaques (centers of infection) on the phage-sensitive control strain. The efficiency of λ*vir* forming centers of infection (ECOI) is calculated as (number centers of infection/ml from test strain) x 100/(number centers of infection/ml from control strain). The ECOI for λ*vir* on P_*lac*_-*dicBF* cells was 3% (Fig. 1C). This result is similar to the results of EOP assays (Fig. 1B), suggesting that the growth characteristics of the P_*lac*_-*dicBF* test strain did not impact the experimental outcome. Deletion of *dicB*, alone or in combination with *dicF*, restored the ECOI to ~80%. The Δ*dicF* strain gave an ECOI of 5% (Fig. 1C). These results are again consistent with our EOP experiments (Fig.1B) implicating DicB as the major player in the phage resistance phenotype.

### The *dicBF* operon promotes resistance against λ, but not other phages

To determine if transient expression of *dicBF* conferred resistance to other phages, we conducted infection experiments using control and P_*lac*_-*dicBF* strains with nine different lytic and temperate phages (Fig. 2). In this experiment, the EOP of λ*vir* on the P_*lac*_-*dicBF* strain was 0.016 or 1.6% compared to the control strain, which was the lowest of the nine phages tested (Fig. 2). Partial resistance was observed for T3 phage, which had an EOP of 0.14 on P_*lac*_-*dicBF* cells. However, the EOP for the remaining seven phages, including phages ϕ80 and HK97, which are closely related to λ, was similar to control cells (Fig. 2). These results suggest DicB does not provide a broad-spectrum of resistance against bacteriophages.

**Figure 2.**
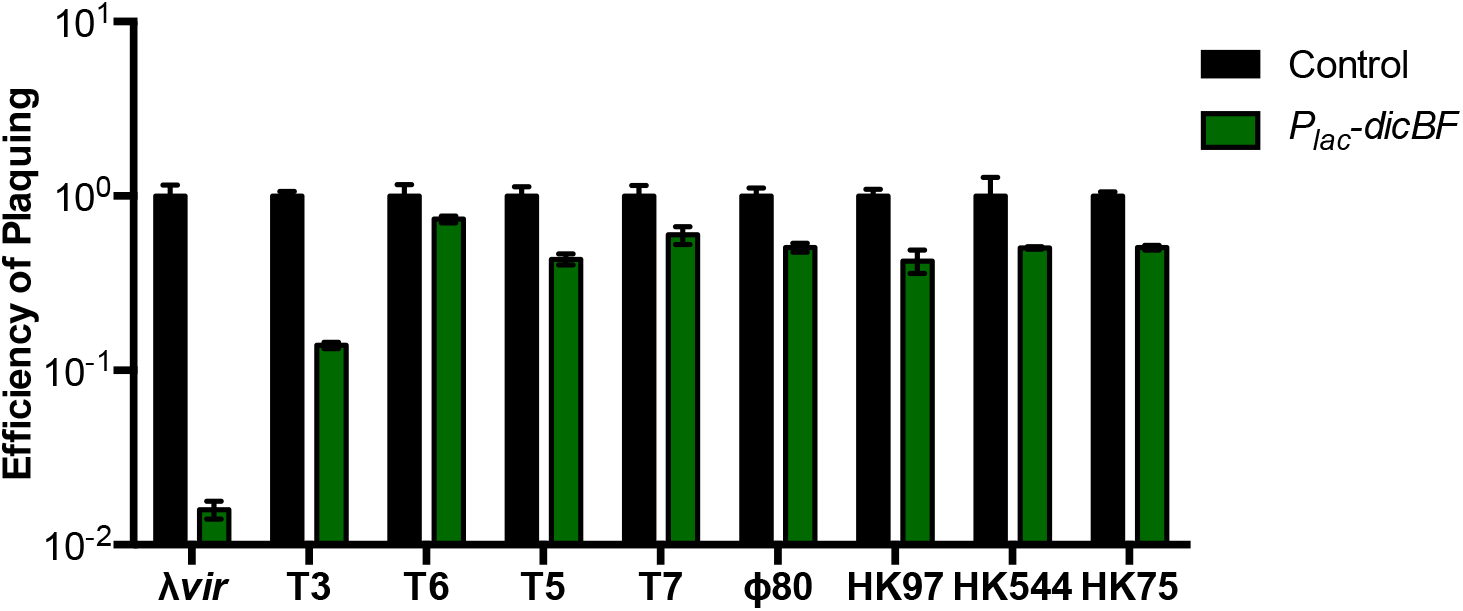
The *dicBF* operon confers resistance against λ phage but not other phages. For each of the nine phages, titer in terms of pfu/ml was calculated by infection of control (DJ480) and P_*lac*_-*dicBF* (DB240) cells. The cells were prepared for infection and the EOP was calculated for each phage as described for Fig.1B. Error bars were calculated as standard deviation of values from three biological replicates.

### Effect of *dicBF* expression on λ phage growth

The classical experiment to study the growth cycle of phage in bacteria is the one-step growth curve, as described by Ellis and Delbrück (43). They observed a latent period where numbers of phage recovered from infected cells remained low as new phage particles were being synthesized inside the host cell. After the latent period is the “burst” where numbers of infectious phage particles increase rapidly as the phage life cycle is completed and cells are lysed to release mature progeny. We conducted one-step growth curves for λ*vir* on control and *dicBF-*expressing strains, essentially as described above for ECOI experiments over a time course following infection. λ*vir* was added at an MOI of 0.1 to control and P_*lac*_-*dicBF* cells resuspended in TM buffer. After phage adsorption, the cells were washed to remove unadsorbed phage and the phage-host complexes were diluted into fresh medium with IPTG (see Materials and Methods). At each time-point, the number of infectious phage particles in each culture was calculated by removing samples and plating for PFU on a phage-sensitive control strain.

As expected based on previous results (Figs. 1B, C), the ECOI for λ*vir* on P_*lac*_-*dicBF* cells was reduced by almost 2 logs compared with the control strain at the early time points, and the reduced numbers of phage produced by P_*lac*_-*dicBF* cells persisted across the phage growth curve (Fig. 3). The latent period for P_*lac*_-*dicBF* cells (75 min.) was ~10 min. longer than for control cells (65 min.) (Fig. 3). The calculated burst sizes were 343 phage/P_*lac*_-*dicBF* cell compared to 169 phage/control cell. This increase in burst size in *dicBF*-expressing cells is likely due to filamentation of cells caused by DicB and DicF. It has been shown before that filamenting cells produce more phage than normal size cells (44, 45). Importantly, we observed that the ~3% of phages that escaped DicB-mediated resistance followed a growth curve similar to that of phages growing on control cells. Collectively, these data led us to hypothesize that DicB affects an early step of the phage life cycle like adsorption or DNA injection, since phages that escape this DicB effect complete a relatively normal life-cycle.

**Figure 3.**
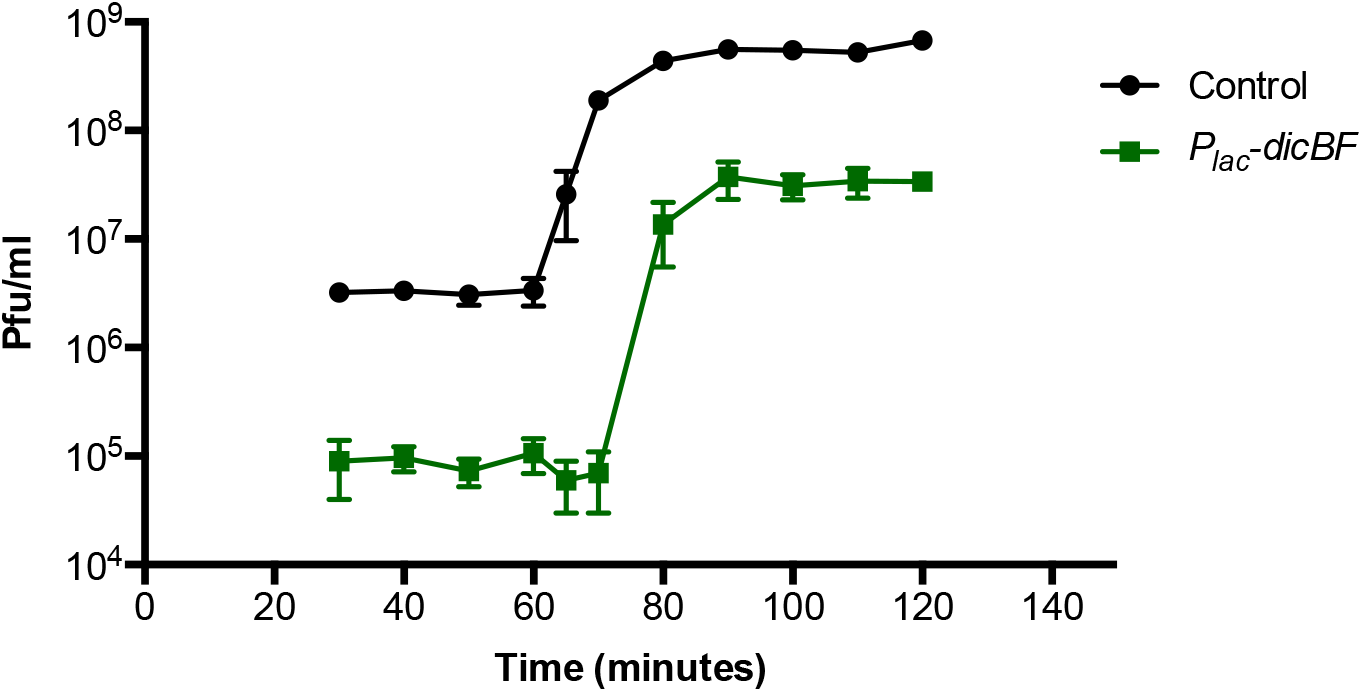
One step growth curve of λ*vir* control and P_*lac*_-*dicBF* cells. The cells were grown with induction of the *dicBF* operon and infection was carried out an MOI of 0.1, similar to the center of infection assay. After removing unadsorbed phages, the cells were diluted in LB medium (with IPTG to induce the *dicBF* operon) and incubated at 37°C for the entire duration of the growth curve. At each time point starting at 30 minutes from the start of infection, samples were removed and added to the phage-sensitive strain (DJ480), and plated to count plaques. Burst size was calculated as (phage titer at 100 minutes - initial titer at 30 minutes)/initial titer at 30 minutes. Latent period was calculated as the time at the mid-point of the exponential phase of the curve. Error bars were calculated as standard deviation of values from three biological replicates.

### The *dicBF* operon does not affect phage adsorption to host cells

To test if expression of the *dicBF* operon affects the first step of phage infection, we tested the ability of λ*vir* to adsorb to host cells expressing this operon. During the ECOI experiment, once phage infection was carried out with cells in TM buffer at an MOI of 0.1 and incubated at 37°C for 10 mins, the phage-cell mixture was centrifuged and the supernatant containing the unadsorbed phages was removed. This supernatant was titered on phage-sensitive control cells by standard plaque assay (residual titer). Control titer was calculated using the same procedure as above, with phage added to TM buffer instead of bacterial cells in the first step of the experiment. The percentage of adsorption was calculated as (control titer – residual titer)*100 / control titer.

The adsorption of λ*vir* to strains expressing the *dicBF* operon was the same as adsorption to the control strain (Table 1). Notably, while λ*vir* and HK97 both adsorb to the same outer membrane receptor, LamB (46, 47), the effect of *dicBF* expression on the EOP of these two phages is significantly different – with reduced EOP only for λ*vir* (Fig. 2). These observations strongly suggest that DicB does not affect the phage life cycle at the step of adsorption to host cells.

**Table 1.** The *dicBF* operon does not affect phage adsorption to host cells. The strains used in this experiment are control (DJ480), P_*lac*_-*dicBF* (DB240), P_*lac*_-*dicBF* Δ*dicB* (DB243), P_*lac*_-*dicBF* Δ*dicF* (DB247) and P_*lac*_-*dicBF* Δ*dicF* Δ*dicB* (DB248). Cells were infected at an MOI of 0.1 with λ*vir* and allowed to adsorb for 10 mins. at 37°C. After adsorption, the samples were centrifuged and the supernatant containing the unadsorbed phages were titered on phage sensitive cells for quantification (residual titer). Control titer was calculated by carrying out the assay with TM buffer, without bacterial cells. The percentage of adsorption was calculated as (control titer-residual titer)*100/ control titer. Standard error was calculated as standard deviation of values from three biological replicates.

| | Control | $P_{lac}-dicBF$ | $P_{lac}-dicBF \Delta dicB$ | $P_{lac}-dicBF \Delta dicF$ | $P_{lac}-dicBF \Delta dicF \Delta dicB$ |
| --- | --- | --- | --- | --- | --- |
| % adsorption | 99 | 99 | 99 | 99 | 99 |
| Standard error (+/-) | 0.22 | 0.25 | 0.69 | 0.23 | 0.15 |

### Recombinant λ phages with the host range region of ϕ80 are not affected by DicB

The genomes of λ phage and ϕ80 (a lambdoid phage) have strikingly similar organization, allowing for easy construction of recombinant phage (48, 49). One prominent difference between λ and ϕ80 is their use of different outer and inner membrane receptors. λ uses LamB (outer membrane) and ManYZ (inner membrane) for adsorption and DNA injection respectively (50–54), while ϕ80 uses FhuA (outer membrane) and the TonB complex (inner membrane) (48, 55, 56). The phage genes encoding determinants for utilization of host outer and inner membrane receptors are located in the host range region of lambdoid phage genomes. Our results so far suggest that the DicB-dependent phage resistance phenotype is not due to an effect on adsorption to the outer membrane receptor but might be mediated at another early step of infection such as injection of the phage genome through the inner membrane receptor. To test this idea, we measured phenotypes of control and *dicBF*-expressing cells challenged with recombinant λ phage containing the host range region of ϕ80 (λh80). The λh80 phage carry most of the wild-type λ genome but have an altered host range region specifying use of the ϕ80 outer and inner membrane receptors. To confirm this, we tested the plaquing ability of λh80 phages on wild-type, Δ*fhuA*, Δ*tonB* and Δ*manXYZ E. coli* strains. As expected, the λh80 phages, like ϕ80, did not plaque on Δ*fhuA* and Δ*tonB* strains but formed normal plaques on the Δ*manXYZ* strain (Table S3). Next, we carried out EOP assays using λ*vir* and a panel of recombinant phage with the host range of λ or ϕ80 (Table S1) on control and P_*lac*_-*dicBF* cells (Fig. 4). We hypothesized that if DicB mediates resistance to λ phage by impairing injection of phage DNA across the cytoplasmic membrane, then phage with the host range of λ would remain inhibited by DicB, whereas λh80 phage with altered inner membrane receptor specificity would not be impacted by DicB. Results of the EOP assays demonstrate that phage with λ host range remain sensitive to DicB-mediated inhibition while λh80 phages had a similar EOP on P_*lac*_-*dicBF* and control cells (Fig. 4). Together with our previous results, this observation suggests that DicB-mediated resistance acts at the level of the inner membrane receptor ManYZ used for λ phage DNA injection into the cytoplasm of *E*. *coli*. We note that the panel of phages that we tested in this experiment had other genetic differences aside from the different host ranges (Table S1). Only the host range was correlated with susceptibility to DicB-mediated resistance.

**Figure 4.**
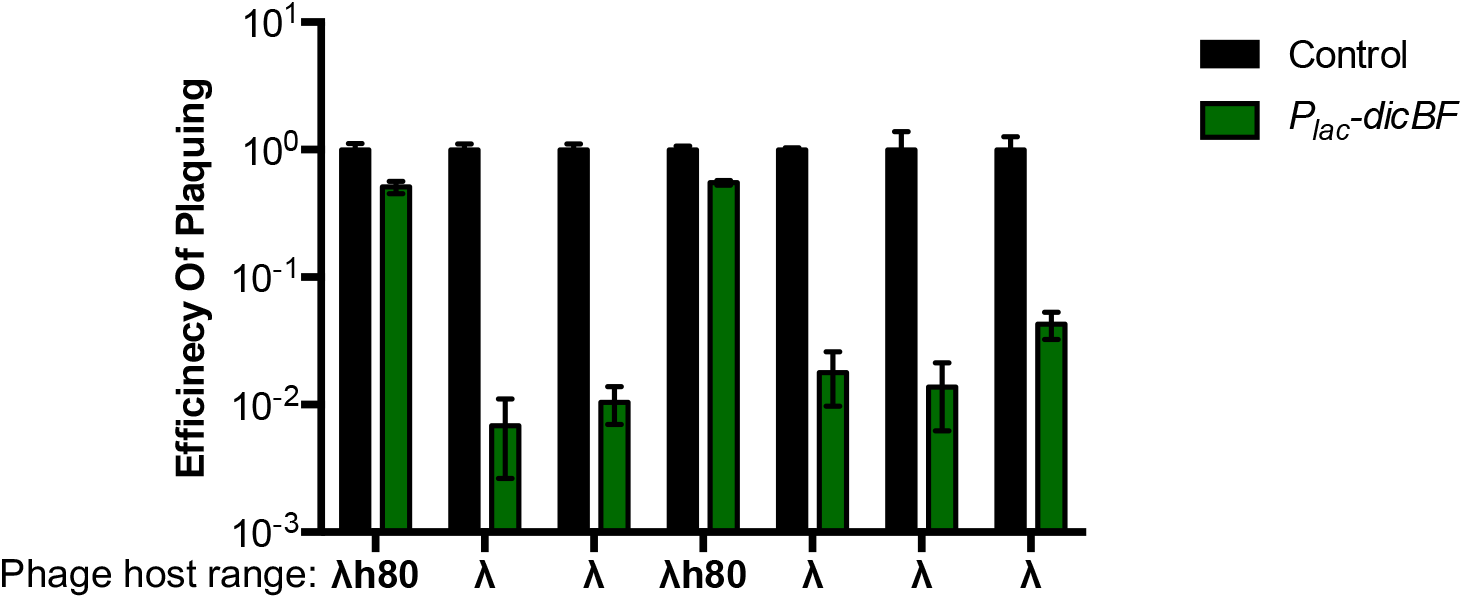
λ phage with the host range of ϕ80 is not affected by DicB. Recombinant λ phages with either the λ or ϕ80 host range were plaqued on control (DJ480) or P_*lac*_-*dicBF* (DB240) cells. Phage names from left to right are: 185, 148, 169, 173, 158, 138 and λ*vir* (see Table S1 for phage genotypes). The cells were prepared for infection and the EOP was calculated for each phage as described for Fig. 1B. Error bars were calculated as standard deviation of values from three biological replicates.

Phage 434 (53) is another phage that uses ManYZ for injection of DNA through the cytoplasmic membrane (Fig. 5A). Previous studies have shown that *manXYZ* deletion mutants (also known as *pel* mutants) were resistant to infection by λ and phage 434, but not ϕ80 (53). To further test our hypothesis that DicB inhibits phage infection at the level of DNA entry through ManYZ, we tested the ability of λ*vir*, phage 434 and ϕ80 to infect control and *dicBF*-expressing cells in *manXYZ*^+^ and Δ*manXYZ* backgrounds. We verified that the λ, phage 434 and ϕ80 phages plaque as expected on wild-type and strains with mutations in specific receptors (Table S4). As shown above (Fig. 2), ϕ80 plaquing efficiency is not impacted by expression of the *dicBF* operon in a *manXYZ*^+^ background. The Δ*manXYZ* mutant host also supported wild-type EOPs for ϕ80 plaquing regardless of whether *dicBF* was expressed (Fig. 5B). For λ*vir*, the EOP on P_*lac*_-*dicBF* cells was 4% relative to control in *manXYZ^+^* cells, whereas the *ΔmanXYZ* host did not support λ*vir* growth (Fig. 5B). The pattern of growth for phage 434 was very similar to that of λ*vir*, with a reduced EOP of ~10% on P_*lac*_-*dicBF* cells in the *manXYZ*^+^ host and no countable plaques on the Δ*manXYZ* host (Fig. 5B). The results of this experiment are consistent with the hypothesis that DicB inhibits the use of the mannose phosphotransferase system proteins, ManYZ, as an inner membrane receptor for productive phage infection.

**Figure 5.**
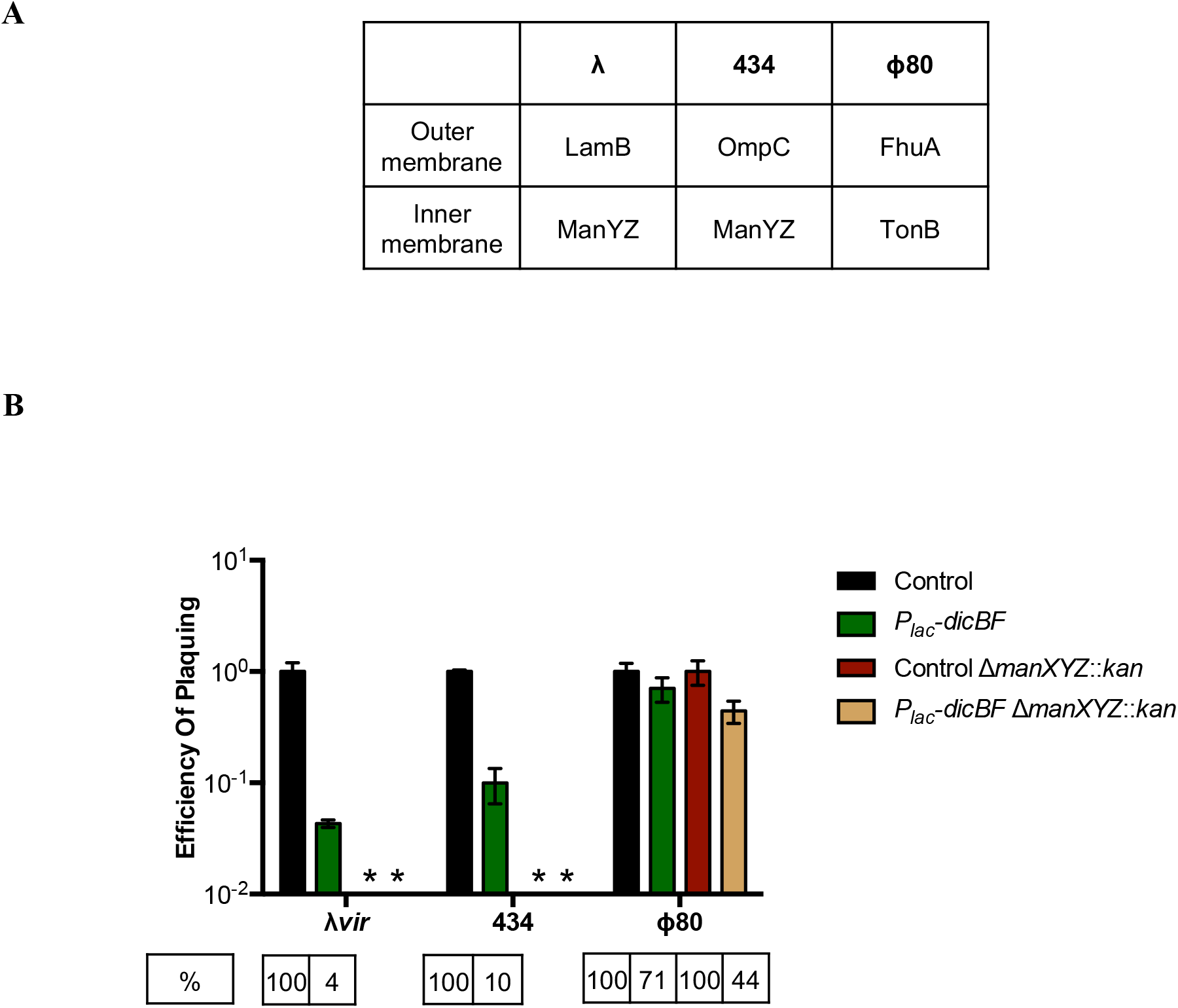
Phage 434 plaquing on ManYZ^+^ strains is inhibited by the *dicBF* operon. (A) Outer and inner membrane receptor specificity of phages λ*vir*, 434 and ϕ80. (B) EOP assay was carried out by preparing cells and calculating titer of phages on the different strains as described in Fig.1B. The strains used in this experiment are control (DJ624), P_*lac*_-*dicBF* (DB240), control Δ*manXYZ*∷kan (PR187) and P_*lac*_-*dicBF* Δ*manXYZ*∷kan (PR191). EOP of phages for each strain is calculated with respect to the control strain in the same background. Error bars were calculated as standard deviation of values from three biological replicates. * denotes strains for which plaques could not be counted.

### Growth of *dicBF*-expressing cells is inhibited on plates with mannose as the C source

Our previous results point to the small protein DicB inhibiting the activity of mannose transporter ManYZ proteins with regard to DNA uptake during phage infection. To test whether DicB inhibits the function of these proteins more broadly, we checked the growth of *dicBF*-expressing cells on mannose as the sole C source. For this experiment, we used control, P_*lac*_-*dicBF*, and P_*lac*_-*dicBF* Δ*dicB* strains. The strains were streaked on M63 minimal plates with different sugars with or without 0.025mM IPTG (to induce *dicBF* expression) and incubated for 44 hours at 37°C (Table 2, Fig. S2). In the absence of inducer, all the strains had near normal growth on the different sugars used. When *dicBF* expression was induced using 0.025mM IPTG, we observed growth inhibition of P_*lac*_-*dicBF* cells on mannose and glucosamine, but not on glucose, fructose or N-acetyl glucosamine. Deletion of *dicB* relieved the growth inhibition on mannose and glucosamine. Both mannose and glucosamine sugars are transported via the ManXYZ transporter in *E. coli* (57–59). These results demonstrate that DicB affects growth specifically on substrates of ManYZ. Growth on sugars that are transported by other PTS proteins was unaffected. These data suggest that DicB impacts at least two different functions of ManYZ – uptake of phage DNA during infection and transport of sugar substrates.

**Table 2.**
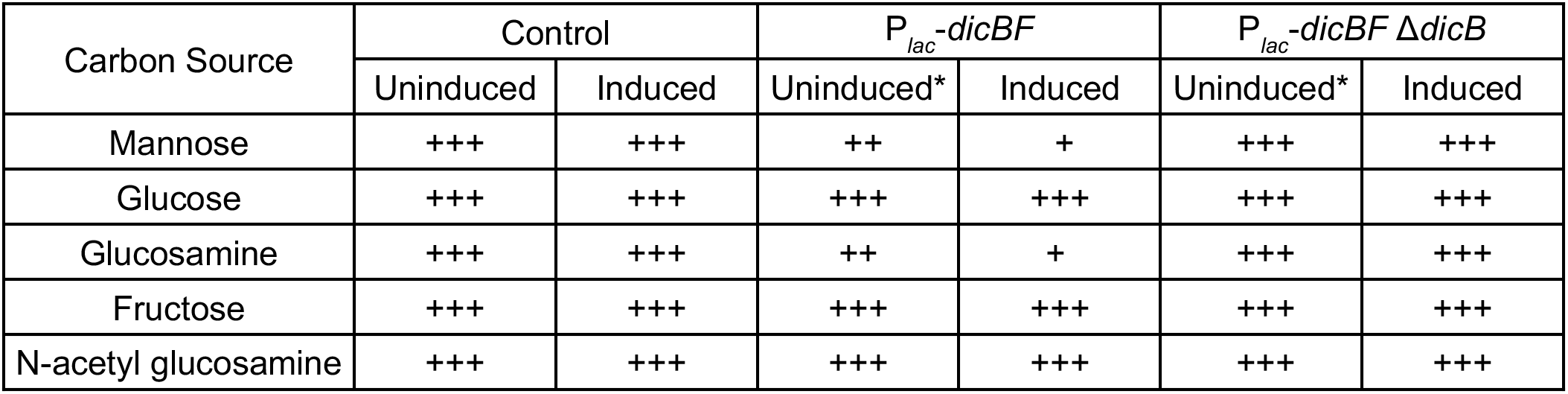
Growth of *dicBF* expressing cells is inhibited on plates with mannose as the C source. The strains were streaked on M63 minimal medium with 0.2% sugars as C source without or with 0.025mM IPTG and incubated for 44 hours at 37°C. The strains used were control (DJ624), P_*lac*_-*dicBF* (DB240) and P_*lac*_-*dicBF* Δ*dicB* (PR165). +++ indicates normal growth, ++ and + indicates decremental growth. * denotes that the P_*lac*_ promoter is leaky and we suspect low level expression of the *dicBF* operon even at 0mM IPTG.

| Carbon Source | Control | | $P_{lac}$-*dicBF* | | $P_{lac}$-*dicBF* Δ*dicB* | |
| --- | --- | --- | --- | --- | --- | --- |
|  | Uninduced | Induced | Uninduced* | Induced | Uninduced* | Induced |
| Mannose | +++ | +++ | ++ | + | +++ | +++ |
| Glucose | +++ | +++ | +++ | +++ | +++ | +++ |
| Glucosamine | +++ | +++ | ++ | + | +++ | +++ |
| Fructose | +++ | +++ | +++ | +++ | +++ | +++ |
| N-acetyl glucosamine | +++ | +++ | +++ | +++ | +++ | +++ |

### MinC mutants that do not interact with DicB lose the phage resistance and sugar phenotypes

The only characterized activity of DicB is inhibition of cell division (20, 60). The mechanism by which DicB impacts cell division requires a protein-protein interaction with MinC, one of the proteins involved in controlling septal ring placement in *E. coli.* MinC is an inhibitor of FtsZ polymerization, and normally MinC concentrations are highest at cell poles so that septum formation is inhibited at polar sites and directed instead to mid-cell (61). Previous work demonstrated that DicB interacts with MinC and brings it to mid-cell via an interaction with ZipA, a septal protein (28). DicB-mediated localization of MinC to cell center inhibits FtsZ polymerization and promotes filamentation (27, 28). To determine if the DicB-MinC interaction is necessary for the DicB-dependent phenotypes we found in this study, we constructed strains with *minC* mutant alleles that produce MinC proteins that are defective for interaction with DicB. We used two different MinC mutants: MinC R172A, which interacts weakly with DicB, and MinC E156A mutant, which does not interact with DicB (37). In strains expressing these *minC* alleles, DicB has a modest (MinC R172A) or no (MinC E156A) impact on cell division, consistent with their reduced binding to DicB. We used MinC E156A and R172A mutant hosts to test whether the DicB-mediated phage resistance or sugar growth phenotypes required the DicB-MinC interaction.

As observed previously, in the wild-type *minC*^+^ background, *dicBF*-expressing cells showed reduced EOP for λ*vir* compared to control cells (Fig. 6). However, in the MinC R172A (reduced binding to DicB) strain, the resistance phenotype was diminished – *dicBF* expression in this host gave an EOP of 12% compared to the control strain. In the MinC E156A (abrogated binding to DicB) background, the EOP of λ*vir* on *dicBF-*expressing cells was very similar to the control strain (Fig. 6). These results suggested that the DicB-MinC interaction is required for the DicB-mediated resistance to λ phage infection. The same strains were grown on M63 minimal plates with different sugars without or with 0.025mM IPTG to induce the *dicBF* operon. As shown above, in the wild-type *minC*^+^ background, expression of *dicBF* inhibited growth on plates with mannose and glucosamine, but not on plates with glucose (Table 3). In contrast, *dicBF* expression in *minC* mutant strains (E156A and R172A) did not inhibit growth on any of the sugars tested (Table 3). Collectively, these data indicate that the new DicB-associated phenotypes we have identified – phage resistance and inhibition of growth on sugars that are transported by ManYZ – require the previously defined molecular mechanism of DicB interaction with the host protein MinC.

**Figure 6.**
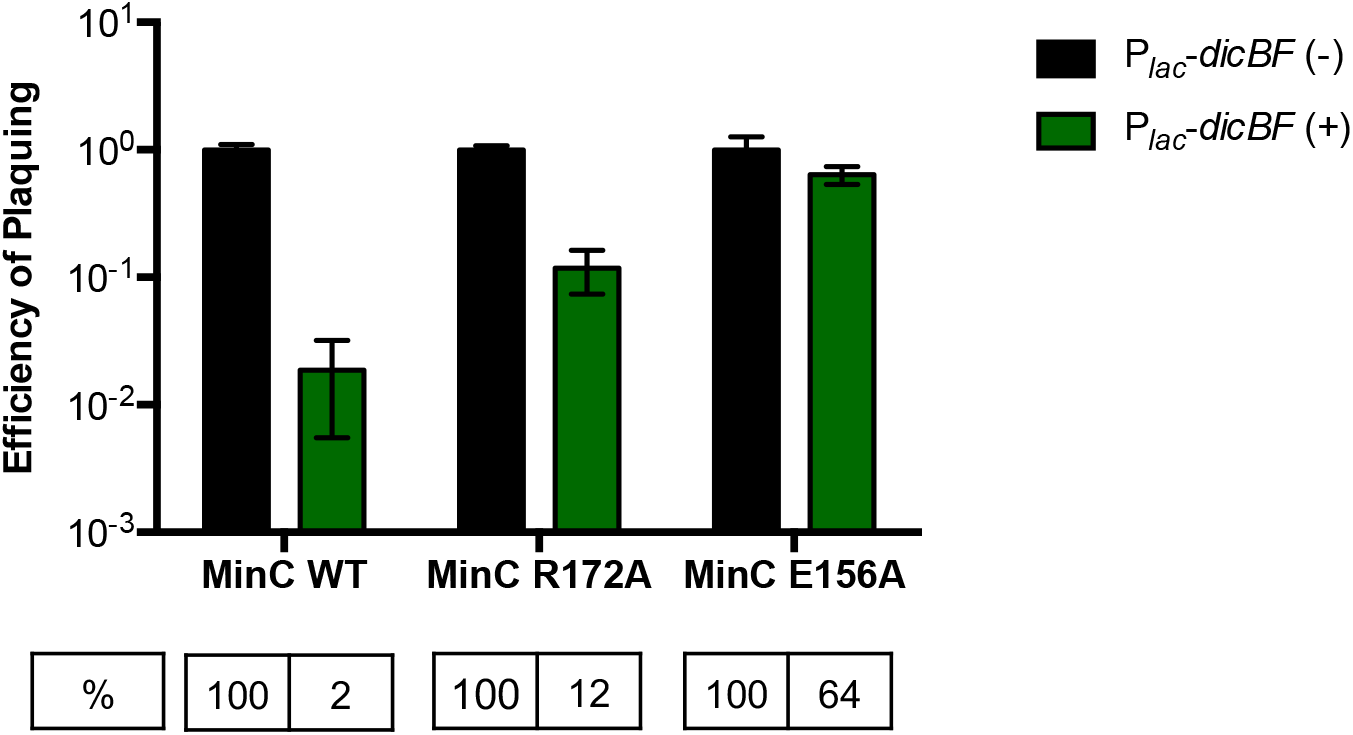
MinC mutants that do not interact with DicB lose the phage resistance effect. The cells were grown with induction of the *dicBF* operon with 0.5mM IPTG, infected with λ*vir* and the EOP calculated as described in Fig.1B. EOP of λ*vir* for each strain is calculated with respect to the control strain in the same background. The strains used in this experiment are control (DJ624), P_*lac*_-*dicBF* (DB240), control *minC* R172A (PR181), P_*lac*_-*dicBF minC* R172A (PR183), control *minC* E156A (PR180) and P_*lac*_-*dicBF minC* E156A (PR182). Error bars were calculated as standard deviation of values from three biological replicates.

**Table 3.**
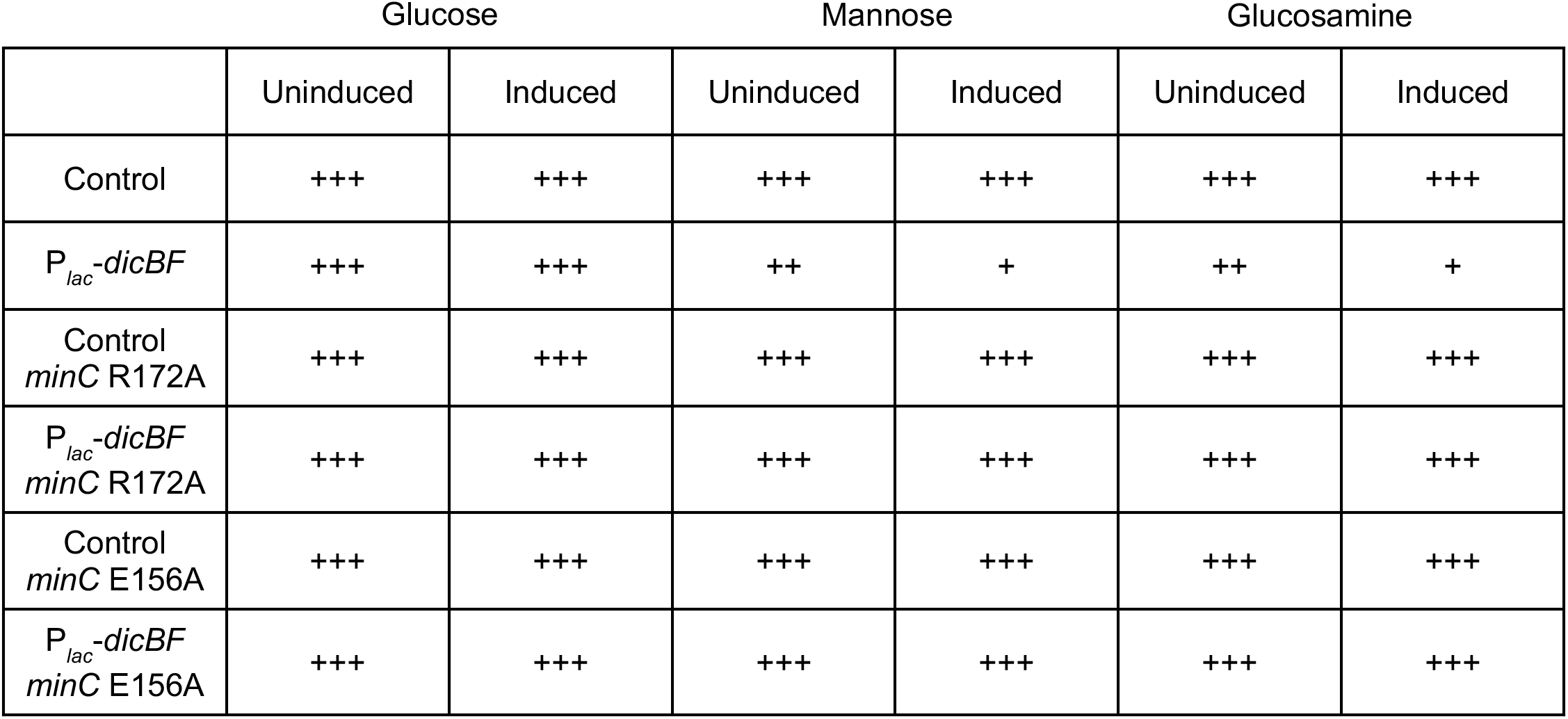
MinC mutants that do not interact with DicB regain ability to grow on Mannose and Glucosamine. The strains were streaked on M63 minimal medium plates with 0.2% sugars and 0.025mM IPTG to induce *dicBF* operon. The plates were incubated for 44 hours at 37°C. The strains used in this experiment are control (DJ624), P_*lac*_-*dicBF* (DB240), control *minC* R172A (PR181), P_*lac*_-*dicBF minC* R172A (PR183), control *minC* E156A (PR180) and P_*lac*_-*dicBF minC* E156A (PR182). +++ indicates normal growth, ++ and + indicates decremental growth. The P_*lac*_ promoter is leaky and we suspect low level expression of the *dicBF* operon even at 0mM IPTG.

## Discussion

The existence of cryptic or defective prophages on bacterial chromosomes was discovered long ago (4), but their potential beneficial functions for host cells are still coming to light. In part, this is because we do not know the functions of the majority of genes encoded on these prophages. In this study, we have identified a new functional role for the cryptic prophage-encoded protein DicB in *E. coli* K12. We showed that induction of the *dicBF* operon makes cells resistant to infection by phages that use the ManYZ PTS proteins as inner membrane receptors for DNA injection (Figs. 1, 5). DicB, a 62-amino acid protein encoded by the *dicBF* operon, plays the primary role in conferring this phage resistance phenotype (Fig. 1). Our results are consistent with the model that DicB inhibits phage DNA injection through the mannose transporter proteins ManYZ (Figs. 4, 5). The DicB effect on ManYZ also inhibits ManYZ-dependent transport of sugar substrates (Tables 2, 3), suggesting that DicB affects the general structure or function of these transport proteins. Previous work demonstrated that DicB inhibits cell division by interacting with and affecting localization and activity of the cell division proteins MinC and FtsZ (23, 26–28). In this study, we found that the DicB-associated phage resistance and sugar utilization phenotypes are dependent on DicB-MinC interactions (Fig. 6).

Prior to this work, the only known function of DicB was inhibition of cell division. DicB directly interacts with MinC of the Min system, which consists of the proteins MinC, MinD and MinE, which play a role in spatial positioning of the FtsZ ring at mid-cell for cell division. MinC is a negative regulator of FtsZ polymerization, and in *E. coli* MinC oscillates between the two cell poles (driven by MinD and MinE) in order to inhibit Z ring assembly at the poles (62, 63). However, when DicB is expressed, a DicB-MinC complex is formed which interacts with the septal ring component ZipA and stimulates Z ring depolymerization at mid-cell leading to cell filamentation (28). Both activities of DicB, cell division inhibition and the ManYZ inhibition phenotypes reported in this study, involve interaction with MinC. These results imply a previously unsuspected link between the Min system or other components of the cell division machinery and the mannose PTS. A few studies have examined localization of various PTS proteins. The general PTS proteins, EI and HPr, were found to localize primarily to cell poles (64, 65). Localization of EII sugar permeases has been less studied, but there is recent evidence that these proteins cluster together around the cell membrane (66). It will be interesting in future work to explore the subcellular localization of ManYZ and examine if or how it is impacted by MinC and DicB.

It is clear that active prophages can protect their hosts from superinfection by other phages (67, 68). One of the earliest identified phage resistance mechanisms is called superinfection exclusion, which can occur by a number of different mechanisms (67, 68). One mechanism of superinfection exclusion is mediated by prophage-encoded proteins that block entry of a superinfecting phage’s DNA by mechanisms that are not well defined, but may involve modifying the activity or function of inner membrane receptors (67). For example, protein gp15 of HK97 prophage inhibits superinfection by HK97 phage by preventing phage DNA entry through the PtsG inner membrane receptor (47, 69). SieA is an inner membrane protein encoded by the P22 prophage of *Salmonella typhimurium* that prevents infection by P22, L, MG178, and MG40 phages by inhibiting DNA entry into the cytoplasm (70, 71). The P1 prophage-encoded Sim protein was also shown to confer superinfection immunity against P1 and few other phages by an unknown mechanism that occurs after adsorption of the infecting phage (72). While these examples illustrate superinfection exclusion by active prophages, our results suggest that in at least some cases defective prophages can play similar roles in protecting their hosts from phage infection.

*E. coli* strains have several defective prophages residing on their chromosomes, many of which have undergone extensive deletions since their original integration (16). There is evidence that many of the genes remaining on the cryptic prophages, including the core phage genes which code for tail and lysis proteins, are under strong purifying selection, implying that these genes now perform functions beneficial to the host (16). *E. coli* K12 has 9 defective prophages, 8 of which do not respond to conditions that usually induce an active prophage (19). Some of these prophages have lost genes important for excision from the host chromosome. For example, CP4-44 prophage entirely lacks an integrase, Qin prophage has an inactive integrase and DLP12 prophage has lost part of the excisionase gene (19, 73). Pathogenic *E. coli* O157:H7 strain Sakai harbors 18 prophages and 17 of these were found to have defects in the prophage genes required for excision and morphogenesis (15). Though not fully functional, three of these 18 defective prophages carry virulence factors and were capable of packaging and transferring their DNA to *E. coli* K12 (15). These observations demonstrate that even though defective prophages have undergone significant genetic changes through their life history with the host, some retain functions that allow them to continue acting as gene-transfer agents.

It has been speculated that another beneficial role of defective prophages could be encoding functions that are important for host cell adaptation to stress conditions. In a study by Wang *et al.* (19), defective prophages of *E. coli* K12 were shown to increase resistance to environmental stresses like oxidative stress, osmotic stress, and to certain antibiotics like quinolones and beta-lactams. This study reported that Δ*qin* derivatives of the parent strain were more sensitive to beta-lactam antibiotics and that Δ*dicB* strains showed greater sensitivity to azlocillin and nalidixic acid (19). We constructed Δ*qin* and Δ*dicB* strains and examined sensitivity to nalidixic acid and ampicillin, and we found no differences in sensitivity between parent strains and mutants (data not shown). It is possible that phenotypes vary with strain background – our strains are MG1655 derivatives and Wang, *et al.*, used BW25113 (19).

Identifying the signals or conditions that induce prophage genes will be key to understanding their physiological roles in host cells. We have exposed our strains to various conditions that are known to induce prophage gene expression, including DNA damage, starvation, and exposure to antibiotics, and have not yet identified conditions that substantially induce transcription from the native *dicBF* promoter (data not shown). Another study (74), reported that *E. coli* K12 MG1655 cells undergo DicF-dependent filamentation under anaerobic conditions (growth in large volume anaerobic fermenters). The authors of this study suggest that stability of DicF is differentially regulated such that it is more stable under anaerobic growth conditions and degraded faster under aerobic conditions. A very recent study found that four DicF orthologs encoded by different prophages in *E. coli* O157:H7 are produced under microaerobic growth conditions (31). These DicF sRNAs promote low oxygen-responsive virulence gene expression via base pairing-mediated regulation of a key virulence transcription factor. These studies suggest that in at least some *E. coli* strain backgrounds, oxygen is an important signal for modulation of *dicBF* operon transcription or DicF mRNA stability. However, we have not observed any DicF- or DicB-mediated filamentation of MG1655 cells grown in small volume LB liquid cultures in an anaerobic chamber (data not shown), so we speculate that additional signals or conditions might contribute to *dicBF* operon expression in our host strain background.

Previous studies from our lab characterized the mRNA target regulon of DicF (25). In addition to the previously discovered DicF target *ftsZ* mRNA, we found that DicF base pairs with and represses translation of *xylR*, *pykA* and *manXYZ* mRNAs, encoding the xylose repressor, pyruvate kinase, and mannose PTS components, respectively (25, 75). Thus, the *dicBF* operon encodes a base pairing-dependent sRNA regulator (DicF) and a small protein (DicB) that act at different levels to inhibit the synthesis and activity of a PTS sugar transporter (ManXYZ). This is strikingly similar to the regulation of the glucose PTS (*ptsG*, enzyme IICB^Glc^) by the dual-function sRNA SgrS and the small protein it encodes, SgrT. SgrS base pairs with and represses translation of *ptsG* mRNA (76, 77), while SgrT inhibits PtsG activity at a post-translational level (78–80). Perhaps regulation of PTS enzyme synthesis and activity by sRNAs and small proteins is a common mechanism for post-transcriptional control of these systems. Future studies on the multitude of sRNAs and small proteins encoded on prophages and bacterial chromosomes promise to reveal more surprising connections between phages and their hosts.

## Acknowledgements

We thank Jeffrey Gardner, Andrei Kuzminov, Alan Davidson, and Sankar Adhya for their generous gifts of various phages which were critical for our experiments. We also thank Nadim Majdalani and John Cronan for providing strains. We are grateful to James Slauch for his help and advice with the design of phage experiments. Finally, we would like to express our thanks to former and current Vanderpool and Slauch lab members for thought-provoking discussions.

This work was supported by the National Institutes of Health (R01 GM092830) and the University of Illinois Department of Microbiology Alice Helm Fellowship to P.T.R.

